# The grass subfamily Pooideae: late Cretaceous origin and climate-driven Cenozoic diversification

**DOI:** 10.1101/462440

## Abstract

**Aim:** Frost is among the most dramatic stresses a plant can experience and complex physiological adaptations are needed to endure long periods of sub-zero temperatures. Due to the need for evolving these complex adaptations, transitioning from tropical to temperate climates is regarded difficult and only half of the world’s seed plant families have temperate representatives. Here, we study the transition from tropical to temperate climates in the grass subfamily Pooideae, which dominates the northern temperate grass floras. Specifically, we investigate the role of climate cooling in diversification.

**Location:** Global, temperate regions

**Time period:** Late Cretaceous-Cenozoic

**Major taxa:** The grass subfamily Pooideae

**Methods:** We date a comprehensive Pooideae phylogeny and test for the impact of paleoclimates on diversification rates. Using ancestral state reconstruction, we investigate if Pooideae ancestors experienced frost and winter. To locate the area of origin of Pooideae we perform biogeographical analyses.

**Results:** We estimated a late Cretaceous origin of the Pooideae (66 million years ago (Mya)), and all major clades had already diversified at the Eocene-Oligocene transition climate cooling (34 Mya). Climate cooling was a probable driving force of Pooideae diversification. Pooideae likely evolved in mountainous regions of southwestern Eurasia in a temperate niche experiencing frost, but not long winters.

**Main conclusion:** Pooideae originated in a temperate niche and experienced cold temperatures and frost long before the expansion of temperate biomes after the Eocene-Oligocene transition. This suggests that the Pooideae ancestor had adaptations to temperate climate and that extant Pooideae grasses share responses to low temperature stress in Pooideae. Throughout the Cenozoic falling temperatures triggered diversification. However, complex mechanisms for enduring strongly seasonal climate with long, cold winters most likely evolved independently in lower taxonomic lineages. Our findings provide insight into how adaptations to historic changes in chill and frost exposure influence distribution of plant diversity today.

## Introduction

Although temperate climates today occupy major parts of the global landmass, they originated and expanded relatively recently in Earth’s history when the global climate started to cool in the late Eocene (Eldrett, Greenwood, Harding, & Huber, 2009; Fine & Ree, 2006; Morley, 2000; Strömberg, 2011; J. Zachos, Pagani, Sloan, Thomas, & Billups, 2001). Many temperate–adapted lineages evolved around and after the Eocene-Oligocene (E-O) transition along with the expansion of cold temperate biomes, especially in the Northern Hemisphere (Favre et al., 2016; Kerkhoff, Moriarty, & Weiser, 2014; Meseguer et al., 2018; Meseguer, Lobo, Ree, Beerling, & Sanmartín, 2015; Near et al., 2012). The concurrence of E-O transition and diversification into temperate climates suggests that the global cooling around 34 Ma ago sparked the evolution of adaptations to cold seasonal climates. Only approximately half of all seed plant families have members in temperate climates (Ricklefs & Renner, 1994). This finding led to the assumption that transitioning from tropical to highly seasonal, cold climates requires the evolution of complex physiological adjustments not so readily accomplished (Donoghue, 2008). Although, (historic) climate cooling likely impacted the evolution of angiosperm lineages, it is still unknown how it influenced today’s distribution of species diversity.

An example of a highly successful temperate lineage is the grass subfamily Pooideae. It is the largest subfamily of Poaceae, comprising almost 4000 species distributed worldwide (Soreng et al., 2017). The remarkable ability to endure in the coldest environments is reflected by its distribution and dominance in temperate and Arctic grass floras (Hartley, 1973; Visser, Clayton, Simpson, Freckleton, & Osborne, 2014), by its capacity for physiological adjustments to tackle the physical damages from cold temperatures and by its precise mechanisms to time life history events such as flowering, growth cessation and cold acclimation to a highly seasonal climate (Fjellheim, Boden, & Trevaskis, 2014; Preston & Sandve, 2013). These capacities are well described in the model grass *Brachypodium distachyon* (tribe Brachypodieae) and in its species-rich sister clade, the “core” Pooideae (Soreng & Davis, 1998). The core Pooideae comprise 3232 species in four tribes (Soreng et al., 2017), and include all commercially important Pooideae crops, like bread wheat (*Triticum aestivum*) and barley (*Hordeum vulgare*) and forage grasses like fescues (*Festuca* spp.) and ryegrass (*Lolium perenne*). The Pooideae share a common ancestor with the tropical and subtropical subfamilies Oryzoideae (previously Ehrhartoideae) and Bambusoideae, and together they form the so-called BOP clade (Soreng et al., 2017). A shift in climatic preferences from warm, tropical to colder, more temperate climates has been inferred in the stem-lineage of the Pooideae (Edwards & Smith, 2010).

The age of Pooideae has, however, been controversial and ages ranging from 45 to 64 Ma have been suggested, depending on the selection of non-Pooideae fossils included in the analyses (Bouchenak-Khelladi, Verboom, Savolainen, & Hodkinson, 2010; Burke, Lin, Wysocki, Clark, & Duvall, 2016; Christin et al., 2014; Prasad et al., 2011; The International Brachypodium Initiative, 2010; Vanneste, Maere, & Van de Peer, 2014; Wang et al., 2015). During the time range of suggested origin for Pooideae the global climate was warm (Mudelsee, Bickert, Lear, & Lohmann, 2014; J. Zachos et al., 2001) and seasonality in temperature relatively low (Archibald, Bossert, Greenwood, & Farrell, 2010). In the late Eocene gradual climate cooling lead to an expansion of temperate climates (Liu et al., 2009; Potts & Behrensmeyer, 1992; J. Zachos et al., 2001). A drop in global temperature around the E-O transition 34 Ma ago (Pound & Salzmann, 2017), followed by increased seasonality (Eldrett et al., 2009; J. Zachos et al., 2001) intensified the expansion of the temperate niche. However, disentangling how falling temperatures throughout the Cenozoic have impacted the evolutionary history of Pooideae is hampered by the lack of a properly dated, comprehensive Pooideae phylogeny.

The controversial age of Pooideae reflects the overall poor fossil record of Poaceae (Strömberg, 2011). Recent findings of old Poaceae fossils firmly reject a Paleogene origin of grasses, and instead suggest a crown age of at least 100 Ma (Poinar Jr., Alderman, & Wunderlich, 2015; Shi et al., 2012; Wu, You, & Li, 2017). Furthermore, 66 Ma old epidermal fragments containing phytoliths (Prasad, Strömberg, Alimohammadian, & Sahni, 2005) showed diagnostic features of subfamily Oryzoideae. Apparently in agreement with an older age for Poaceae, recent dating studies have indicated an older age also for Pooideae (Christin et al., 2014; Marcussen et al., 2014; Prasad et al., 2011). These age estimates were however largely based on calibrations external to Pooideae and may not imply accuracy for inferred Pooideae ages in the presence of rate heterogeneity.

In this study we aim to reconstruct the paleoclimatic impact on phylogenetic and diversification history of the grass subfamily Pooideae. Firstly, we provide a comprehensive, fossil-dated chloroplast phylogeny of the Pooideae subfamily. Fossilisation rates are low in Poaceae and therefore ages are likely to be underestimated using classical node dating where the oldest available fossil is applied as a minimum age constraint. We circumvent this problem by employing a new method (PyRate; Silvestro, Schnitzler, Liow, Antonelli, & Salamin, 2014) that estimates the speciation time probability distribution based on the entire fossil record for each clade, thereby also eliminating the subjective choice of a maximum age constraint. Secondly, we estimate diversification rates and test for an impact of paleoclimates on diversification trajectories, using paleoenvironmental birth-death models. Thirdly, we reconstruct the temperature niche of the Pooideae to establish if the ancestor experienced frost or longer periods of cold temperatures. Lastly, we reconstruct the biogeographical history of Pooideae using ancestral state reconstruction (ASR) methods.

## Materials and Methods

### Materials and sampling

A data matrix containing three cpDNA regions (*matK*, *ndhF*, *rbcL*) for 396 species, (including eight outgroup species from Bambusoideae, Oryzoideae, Panicoideae), was assembled from GenBank and from own accessions (see Table S1.1 in supporting information). Sampling was aimed at being exhaustive at the genus and tribe levels. Sequences for some taxa were obtained *de novo* by PCR and Sanger sequencing using custom Pooideae-specific primers (Table S1.2). Clustal alignments were generated and manual adjusted in BioEdit (Hall, 1999). The final alignments for *matK*, *ndhF* and *rbcL* were 1605, 899 and 717 nucleotides long, respectively (sum: 3221), and had 2% (8), 14% (54) and 34% (135) of missing sequences, respectively (average: 17%).

### Sampling of fossils and estimation of fossil origination times

Based on the Pooideae fossil record (Iles, Smith, Gandolfo, & Graham, 2015; Strömberg & McInerney, 2011; Thomasson, 1988) we identified six calibration points (see Appendix 2 for justification of fossils used) and estimated the probability density of origination time for the respecitve clade (Table 1) using PyRate (Silvestro et al., 2014). In short, a fossil matrix, containing associated lineage minimum and maximum age bounds, was entered in PyRate and run for 1 million generations. Ages were sampled randomly from within the age bounds under a uniform probability distribution. The analyses were replicated 20 times. For each lineage, the posterior distribution of speciation times was summarised over all replicates in Tracer v1.6.0 (Rambaut, Suchard, & Drummond, 2013). Mean, variance, 0.025 and 0.975 quantile (Table 1) were used to approximate speciation times to log-normal distributions using ParameterSolver v3.0 (Cook, Wathen, & Nguyen, 2013), which were then used as calibration priors in a conventional BEAST dating analysis.

**Table 1.** Calibrations used for the dating analysis of Pooideae including secondary calibrations derived from an initial dating analysis of Poaceae.

|  |  | Posterior from PyRate |  |  |  | Calibration priors for Poaceae BEAST analysis |  |  |  | Posterior from Poaceae BEAST analysis |  |  | Calibration priors for Pooideae BEAST analysis |  |  |  |
| --- | --- | --- | --- | --- | --- | --- | --- | --- | --- | --- | --- | --- | --- | --- | --- | --- |
| Node | Fossil(s) | mean | variance | 0.025Q | 0.975Q | type | offset | mean | stdev | mean | 0.025Q | 0.975Q | type | offset | mean | stdev |
| <u>Pooideae:</u> |  |  |  |  |  |  |  |  |  |  |  |  |  |  |  |  |
| *Hesperostipa* stem | *Berriochloa* spp. | 30.746 | 11.740 | 26.022 | 37.377 | → logN | 26 | 1.348 | 0.648 | . | . | . | → logN | 26 | 1.348 | 0.648 |
| narrow-leaved Loliineae stem | *Festuca* cf. *capillata* | 27.352 | 96.976 | 16.252 | 45.084 | → logN | 16 | 2.149 | 0.749 | . | . | . | → logN | 16 | 2.149 | 0.749 |
| *Lygeum* stem | *Lygeum* sp. | 19.945 | 10.441 | 16.000 | 26.302 | exp | 16 | 50 | . | . | . | . | exp | 16 | 50 | . |
| *Nassella* stem | *Nassella* spp. | 17.590 | 67.934 | 8.020 | 34.234 | → logN | 8 | 1.984 | 0.744 | . | . | . | → logN | 8 | 1.984 | 0.744 |
| *Piptatheropsis* stem | *Paleoeriocoma hitchcockii* | 17.166 | 54.198 | 8.053 | 31.864 | → logN | 8 | 1.967 | 0.706 | . | . | . | → logN | 8 | 1.967 | 0.706 |
| Stipeae stem | *Stipa florissantii* | . | . | . | . | exp | 34 | 50 | . | . | . | . | exp | 34 | 50 | . |
| Pooideae stem | . | . | . | . | . | . | . | . | . | 71.0 | 60.6 | 80.9 | → N | . | 71.0 | 4.5 |
| BOP crown | . | . | . | . | . | . | . | . | . | 76.0 | 68.9 | 83.4 | → N | . | 76.0 | 3.5 |
| <u>non-Pooideae Poaceae:</u> |  |  |  |  |  |  |  |  |  |  |  |  |  |  |  |  |
| Poaceae stem | Palaeoclaviceps parasiticus (ergot) | . | . | . | . | exp | 98.8 | 2 | . | . | . | . | . | . | . | . |
| Oryzeae stem | *Changii indicum* | . | . | . | . | exp | 66 | 10 | . | . | . | . | . | . | . | . |
| Chusquea stem | *Bambusoideae* cf. *Chusquea* | . | . | . | . | exp | 33.9 | 10 | . | . | . | . | . | . | . | . |
| *Leersia* stem | *Leersia seifhennersdorfensis* | . | . | . | . | exp | 30.7 | 10 | . | . | . | . | . | . | . | . |

→, arrows link posterior estimates in initial analysis with priors in the Pooideae dating analysis; exp, exponential distribution; N, normal distribution; logN, log-normal distribution; offset, oldest possible fossil age, mostly inferred stratigraphically.

### Dating Pooideae

To obtain age priors for the two deepest nodes of Pooideae, we first performed a separate dating analysis in BEAST of a small Poaceae dataset, consisting of cpDNA (*matK*, *ndhF*, *rbcL*) sequences for 41 taxa sampled from all the major Poaceae lineages, and eight fossil constraints (Appendix 2). Resulting ages (Table 1) for the BOP crown and Pooideae stem node were used as normal-distributed priors in the Pooideae dating analysis.

The dating analysis was set up in BEAUti v1.7.4 and perfomred in BEAST v1.8 (Drummond & Rambaut, 2007; Drummond, Suchard, Xie, & Rambaut, 2012). The three cpDNA partitions were analysed using unlinked site models, linked clocks, and linked partition trees. Nucleotide substitution model priors were set to GTR + G (four gamma categories) for all partitions, as suggested by JModelTest v.2.1.10 (Darriba, Taboada, Doallo, & Posada, 2012). To account for rate heterogeneity among lineages the tree was given an uncorrelated lognormal relaxed molecular clock prior assuming a Yule speciation process (birth-only). Substitution rates were given an uninformative uniform prior between 0 and 0.01. For the three most basal nodes topology was constrained by enforcing monophyly for Pooideae, Pooideae + Bambusoideae, and the BOP clade. Constraintas for fossil and secondary calibrations are shown in Table 1. Two MCMC chains were run for 200 million generations while parameters were logged every 20,000 generations. After confirming proper chain mixing, convergence (i.e. ESS >200) and burn-in removal(20 million generations) the two chains were merged in LogCombiner v1.7.4 (part of BEAST package), We used TreeAnnotator v1.7.4 (part of BEAST package) to summarise the data in a maximum credibility (MC) tree with mean node heights

### Dating Poaceae and monocots using new fossils

Recently, three fossils of grasses or putative grasses (Poinar Jr. et al., 2015; Prasad et al., 2011; Wu et al., 2017) have been discovered indicating an older age of Poaceae than previously assumed. To estimate the impact of these fossils on the age of monocots and Poaceae, we performed an additional set of dating analyses on an independent 4-locus dataset for 46 angiosperm species (Table S1.3), including 29 grasses, nine non-grass monocots and one outgroup. Sequences for seven markers, i.e. *atpA* from the mitochondrome (mtDNA), *atpB*, *matK*, *ndhF* and *rbcL* from cpDNA, and *phyB* and *topo6* from the nuclear genome (nDNA), were downloaded from GenBank and aligned with Clustal. For node dating we identified 11 angiosperm fossils (Table S1.4; Barba-Montoya, dos Reis, Schneider, Donoghue, & Yang, 2018; Iles et al., 2015). Additionally, we constrained angiosperm crown node with an age prior (~144 Ma) obtained by a PyRate analysis of the entire angiosperm fossil record (Silvestro, Cascales-Miñana, Bacon, & Antonelli, 2015); to our knowledge, this is the most objective estimate for this node published to date. Dating analyses were then run with and without the two “new” fossils. The taxon sampling necessitated the use of a secondary calibration on the *Brachypodium* stem lineage (age from Table 2) for the former analysis. The dating analysis was performed as described above, except for using an unlinked site model for the four DNA partitions cpDNA, *atpA*, *phyB* and *topo6*, using the HKY + G nucleotide subtitution model for the latter three partitions and running the MCMC chains for 100 million generations (parameters were logged every 10,000 generations). Used node age constraints are given in Table 1.

**Table 2.**
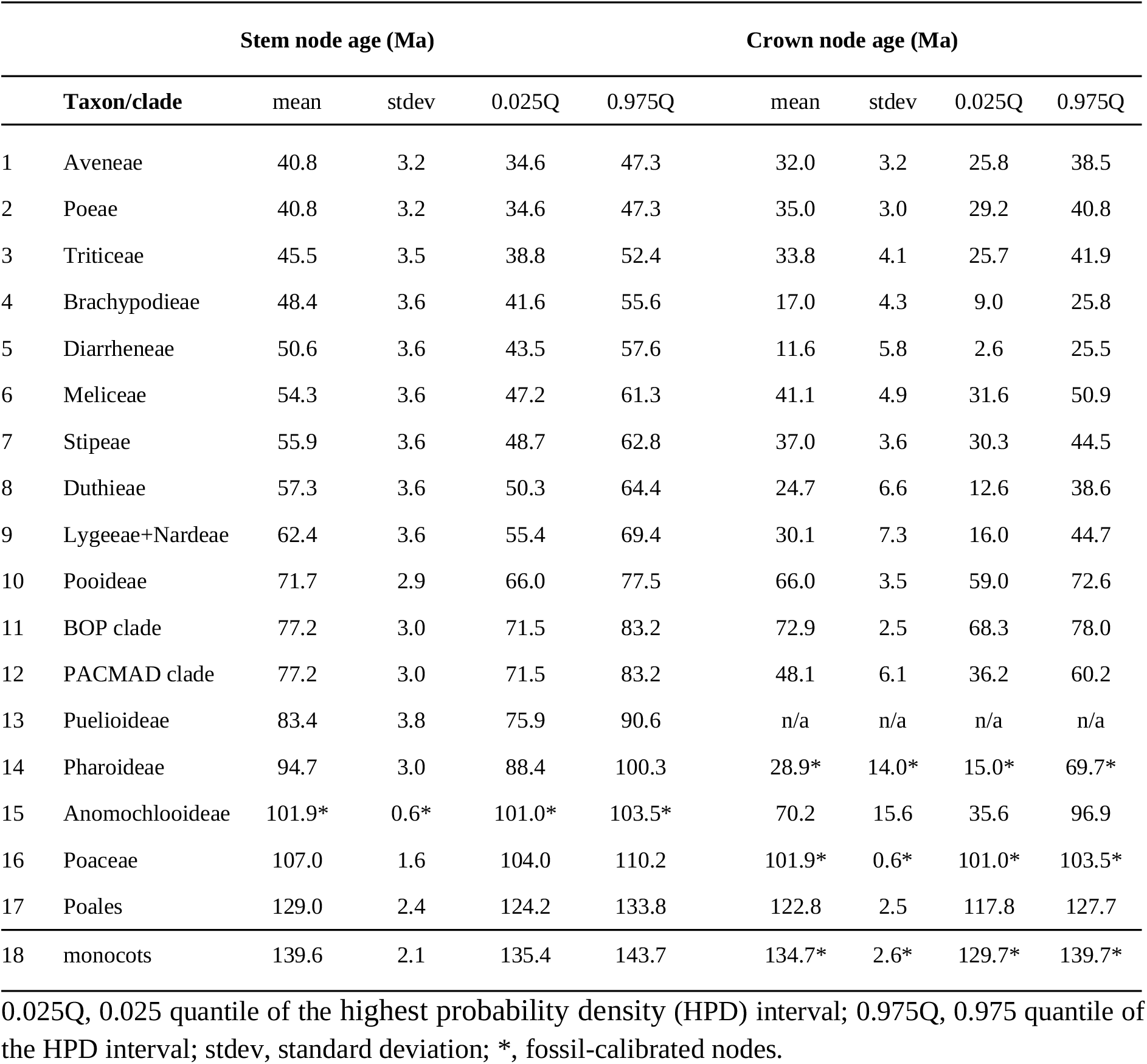
Estimated node ages for Pooideae (rows 1-10) and other monocot clades (rows 11-18) obtained by fossil-calibrated dating analyses.

|  |  | Stem node age (Ma) |  |  |  | Crown node age (Ma) |  |  |  |
| --- | --- | --- | --- | --- | --- | --- | --- | --- | --- |
|  | Taxon/clade | mean | stdev | 0.025Q | 0.975Q | mean | stdev | 0.025Q | 0.975Q |
| 1 | Aveneae | 40.8 | 3.2 | 34.6 | 47.3 | 32.0 | 3.2 | 25.8 | 38.5 |
| 2 | Poeae | 40.8 | 3.2 | 34.6 | 47.3 | 35.0 | 3.0 | 29.2 | 40.8 |
| 3 | Triticeae | 45.5 | 3.5 | 38.8 | 52.4 | 33.8 | 4.1 | 25.7 | 41.9 |
| 4 | Brachypodieae | 48.4 | 3.6 | 41.6 | 55.6 | 17.0 | 4.3 | 9.0 | 25.8 |
| 5 | Diarrheneae | 50.6 | 3.6 | 43.5 | 57.6 | 11.6 | 5.8 | 2.6 | 25.5 |
| 6 | Meliceae | 54.3 | 3.6 | 47.2 | 61.3 | 41.1 | 4.9 | 31.6 | 50.9 |
| 7 | Stipeae | 55.9 | 3.6 | 48.7 | 62.8 | 37.0 | 3.6 | 30.3 | 44.5 |
| 8 | Duthieae | 57.3 | 3.6 | 50.3 | 64.4 | 24.7 | 6.6 | 12.6 | 38.6 |
| 9 | Lygeae+Nardeae | 62.4 | 3.6 | 55.4 | 69.4 | 30.1 | 7.3 | 16.0 | 44.7 |
| 10 | Pooideae | 71.7 | 2.9 | 66.0 | 77.5 | 66.0 | 3.5 | 59.0 | 72.6 |
| 11 | BOP clade | 77.2 | 3.0 | 71.5 | 83.2 | 72.9 | 2.5 | 68.3 | 78.0 |
| 12 | PACMAD clade | 77.2 | 3.0 | 71.5 | 83.2 | 48.1 | 6.1 | 36.2 | 60.2 |
| 13 | Puelioideae | 83.4 | 3.8 | 75.9 | 90.6 | n/a | n/a | n/a | n/a |
| 14 | Pharoideae | 94.7 | 3.0 | 88.4 | 100.3 | 28.9* | 14.0* | 15.0* | 69.7* |
| 15 | Anomochlooideae | 101.9* | 0.6* | 101.0* | 103.5* | 70.2 | 15.6 | 35.6 | 96.9 |
| 16 | Poaceae | 107.0 | 1.6 | 104.0 | 110.2 | 101.9* | 0.6* | 101.0* | 103.5* |
| 17 | Poales | 129.0 | 2.4 | 124.2 | 133.8 | 122.8 | 2.5 | 117.8 | 127.7 |
| 18 | monocots | 139.6 | 2.1 | 135.4 | 143.7 | 134.7* | 2.6* | 129.7* | 139.7* |

0.025Q, 0.025 quantile of the highest probability density (HPD) interval; 0.975Q, 0.975 quantile of the HPD interval; stdev, standard deviation; \*, fossil-calibrated nodes.

### Diversification analyses

We detected changes in diversification rates at estimated points in time using episodic birth-death models (Table 3) in TreePar 2.1 (Stadler, 2011). Potential rate shift times were evaluated in a grid of 1 Ma time intervals. The sampling fraction for the first time interval (present) was set to 0.1 according to the taxon sampling in the present, while the probability of survival per lineage was alternatively set to 1 (no extinction, *i.e.* the sampling of deep branches is complete) or 0.1 (10% of lineages survive to the next period). The likelihood ratio test was used to compare nested models of increasing complexity, from 1 to 8 additional rate shifts.

**Table 3.**
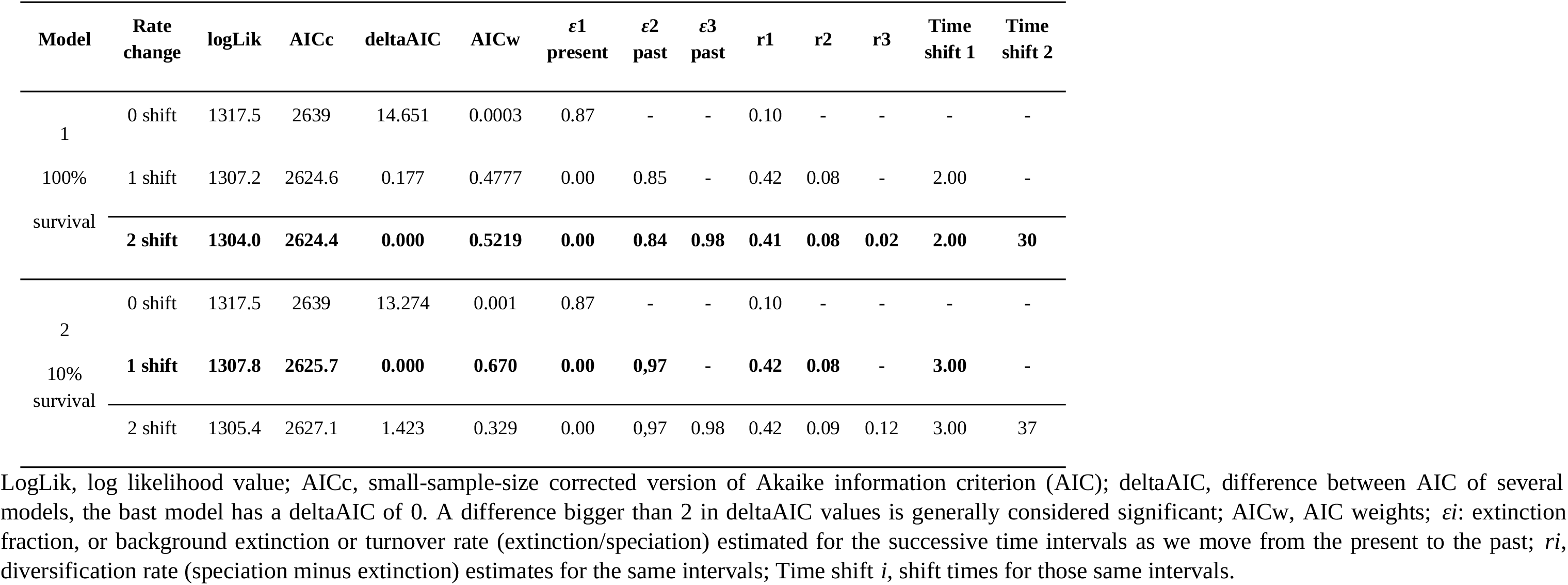
Results for diversification analyses in TreePar for two rate shifts in two different models.

We tested for the impact of paleoclimates on diversification using paleoenvironmental birth-death models (Condamine, Rolland, & Morlon, 2013) implemented in RPANDA (Morlon et al., 2016). In these models, speciation (λ) and extinction rates (μ) can change according to an environmental variable, which itself varys through time. Global paleotemperature data were retrieved from Condamine et al. (2013) and based on the global Cenozoic deep-sea oxygen isotope record as a proxy for global temperatures (J. C. Zachos, Dickens, & Zeebe, 2008). We accounted for incomplete taxon sampling and compared the fit of a set of models where diversification rates vary continuously as a function of an environmental variable (temperature) against a constant-rate birth–death model and a set of models in which speciation and/or extinction vary continuously according to time (Table 4). When rates varied with temperature or time, we assumed exponential variation. For example, if the parameter controlling the variation of λ with time (α) is estimated to be positive, it means that the speciation rate decreases exponentially from the past to the present in association with decreasing global temperatures; if the parameter controlling the variation in μ with time (β) is estimated as negative, extinction increases with decreasing temperatures. With other words, positive values mean a positive effect of the environment on speciation or extinction.

**Table 4.** Results for the paleoenvironment-dependent diversification model in RPANDA.

| Models | #p | logL | AICc | deltaAIC | AICw | $\lambda$ | $\alpha$ | $\mu$ | $\beta$ |
| --- | --- | --- | --- | --- | --- | --- | --- | --- | --- |
| **Birth.cst-Death.cst** | 2 | -1317.53 | 2639.09 | 9.04 | 0.01 | 0.81 | NA | 0.71 | NA |
| **Birth. Time.var-Death.cst** | 3 | -1326.54 | 2659.15 | 33.18 | 0.00 | 0.35 | -0.05 | 0.00 | NA |
| **Birth.cst-Death. Time.var** | 3 | -1314.75 | 2635.55 | 15.83 | 0.00 | 0.72 | NA | 0.59 | 0.00 |
| **Birth. Time.var-Death- Time.var** | 4 | -1314.18 | 2636.46 | 21.01 | 0.00 | 0.68 | -0.02 | 0.51 | -0.02 |
| **Birth.Env.var-Death.cst** | 3 | **-1313.01** | **2632.08** | **0.000** | **0.15** | **0.69** | **-0.22** | **0.00** | **NA** |
| **Birth.cst-Death.Env.var** | 3 | -1314.23 | 2634.52 | 8.566 | 0.01 | 0.68 | NA | 0.51 | 0.02 |
| **Birth.Env.var-Death-Env.var** | 4 | -1312.67 | 2633.45 | 11.66 | 0.00 | 0.70 | -0.16 | 0.18 | -0.10 |

Birth.cst-Death.cst, birth-death model with constant diversification rates; Birth. Time.var-Death.cst, speciation rate varies through time, while extinction rate remains constant; Birth.cst-Death. Time.var, speciation rate remains constant, while extinction rate varies through time; Birth. Time.var-Death- Time.var, both speciation and extinction rate vary through time; Birth.Env.var-Death.cst, speciation rate varies according to temperature, while extinction remains constant; Birth.cst-Death.Env.var, extinction varies with temperature but speciation remains constant; Birth.Env.var-Death.Env.Var, both speciation and extinction rates vary according to the temperature; #p, the number of parameters; logL, log likelihood; AICc, deltaAICc and AICw equivalent to table 3; $\lambda$ , value for speciation rate; $\mu$ , value for extinction rate; $\alpha$ and $\beta$ , correlation parameters between time or paleoenvironmental (temperature) data and speciation and extinction rates, respectively.

### Geo-referenced records and climatic data

Geo-referenced records for Pooideae and outgroup taxa were downloaded from the Global Biodiversity Information Facility (GBIF; www.gbif.org) using rgbif package in R (Chamberlain, 2017). To exclude unreliable records we discarded coordinates with less than three decimals and employed additional filtering implemented in the SpeciesGeoCoder package (Töpel et al., 2016) in R. Taxa with synonymous names were merged using taxonomic information from GBIF. For each of the filtered geo-referenced records, 19 Bioclim variables were downloaded from the WorldClim database (http://www.worldclim.org/) in a 2.5 arc-minutes resolution using the ‘raster’ package in R. After excluding the lower and upper 5% of each Bioclim variable and taxon we calculated mean and standard deviation, which were used in downstream phylogenetic analyses.

### ASR of climatic space

To assess the phylogenetic information contained in each Bioclim variable we determined the phylogenetic signal represented by Pagel’s lambda (λ) (Pagel, 1999) using phylosig function of the R package phytools (Revell, 2012). For the Bioclim variable with the strongest phyolgenetic signal (BIO3, isothermality) we reconstructed ancestral states as continuous trait. To investigate the Pooideae most recent common ancestor’s (MRCA) exposure to frost and prolonged cold, we scored BIO6 (minimum temperature of the coldest month) and BIO11 (mean temperature of the coldest quarter) as binary traits. In case of BIO6, we scored a single binary trait, measuring exposure to frost as either “yes” (BIO6 <0) or “no” (BIO6 ≥0). To measure adaptation to winter severity, we explored which temperatures of BIO11 comprised the strongest phylogenetic signal. We found that the number of months below a mean BIO11 temperature of 2°C and 3°C exhibited the two strongest phylogenetic signals (data not shown). Based on these findings we scored BIO11 as three binary traits, whether BIO11 was below (“yes”), or not below (“no”) 2 °C, 3 °C and 4 °C, respectively. To test if the binary traits were distributed non-randomly along the phylogeny we estimated Fritz & Purvis’ (2010) *D* using the phylo.d function in the R package caper (Orme, 2013; Table S1.5). The continuous and binary ASRs were performed with BEAST using a separate data partition on the previously obtained timetree. We supplied the MC timetree as starting tree and turned off all tree operators. The respective trait was assumed to evolve under a symmetrical Brownian motion (continuous ASR) or an asymmetrical discrete model (binary ASR), and under the same clock as the nucleotide partitions. Remaining settings and priors were left unchanged.

### Historical biogeography

For the discrete biogeographic analysis, each taxon was scored as native to either of six broadly defined biogeographic regions based on GBIF distribution data. A total of 82 out of the 396 taxa were present in more than one biogeographic region, or spanning adjacent regions, and were therefore given trait state “missing”. A discrete ASR analysis was run in BEAST as described above, except the biogeographic trait was assumed to evolve under a symmetrical model.

The discrete analysis identified Eurasia as the ancestral area of Pooideae. To obtain a better resolution of the geographical origin, we performed a continuous analysis for native Eurasian taxa. For each taxon we determined the medians for longitude and latitude as continuous traits, and treated both traits as separate variables in a single ASR analysis. Because taxa distributed outside of Eurasia had arrived there by long-distance dispersal, hence violating the model assumption of Brownian motion, these were given trait state “missing”.

## Results

### Dating analysis

We calculated the origination time of six Pooideae clades (Table 1) to calibrate the corresponding nodes of the phylogeny. The 396-species fossil-calibrated chloroplast phylogeny (Fig. 1a) gave high branch support, i.e. posterior probability (*pp*) = 1, for all the main Pooideae lineages, however lower support was observed for internal branches, owing to missing sequence data. The relative placement of the Meliceae and Stipeae tribes remained unresolved (Fig. 1a). The ages of key nodes are given in Table 2. The crown node of Pooideae (stem node of Brachyelytreae) was inferred to a 95% credibility interval (CI) of 59– 72 Ma (mean 66 Ma), at the Cretaceous-Paleocene boundary (Fig. 1). Reconstructed ages for the stem nodes of the main subclades of Pooideae showed a rapid succession of speciation events between 55–69 Ma (mean 62 Ma; Lygeeae+Nardeae stem) and 44–58 Ma (mean 51 Ma; Diarrheneae stem). The stem node of Brachypodieae, i.e. the split between *Brachypodium* and core Pooideae, was reconstructed to 42–55 Ma (mean 48 Ma). The 95% CI of the crown node ages for the largest subclades, i.e. Stipeae, Meliceae and the three core Pooideae lineages, all overlapped with the E-O transition boundary 34 Ma ago (means 32–41 Ma, 95% CI 26–51 Ma) (Figs. 1a,b; Table 2).

**Figure 1.**
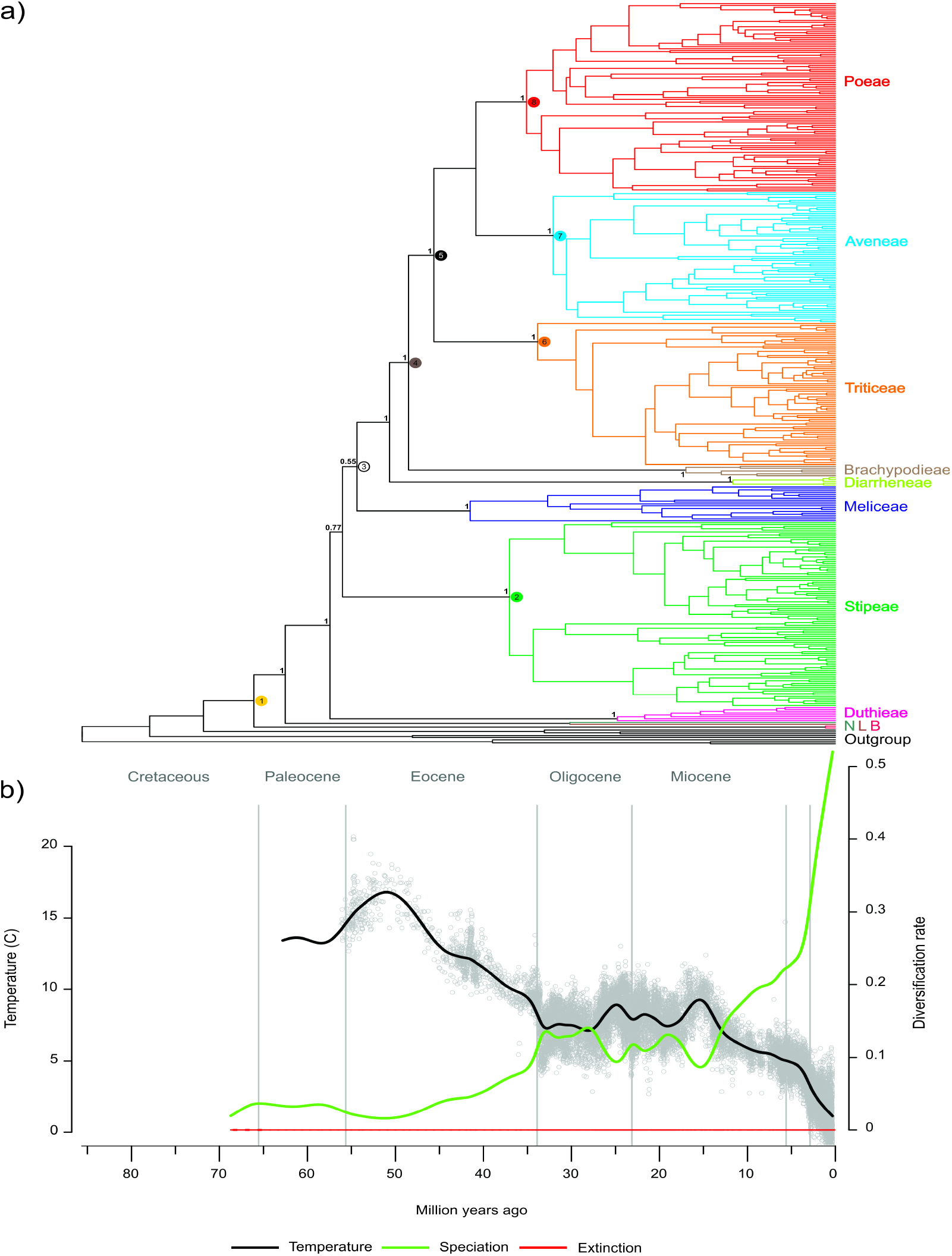
Fossil calibrated Pooideae phylogeny and historic diversication rates. a) Time-tree for Pooideae based on cpDNA sequences (*matK*, *ndhF*, *rbcL*) for 396 species, including eight outgroups, fossil-calibrated at six internal Pooideae nodes. Pooideae tribes are colour-coded and indicated with name. Numbered nodes refer to inferred, historical coordinates of the continuous biogeographical analysis (Fig. S3.3). b) the black line represents the global mean temperature curve from the Paleocene to the present, modified from Zachos *et al.*, (2008). Paleotemperature-dependent diversification model: extinction rates (in red) in Pooideae remain low and constant through time, while speciation rates (in green) increase with the decreasing global temperatures.

### Dating analyses for Poaceae and monocots

To elucidate how our age estimates for Pooideae and recently published grass fossils (Prasad et al., 2011; Wu et al., 2017) might affect age estimates for Poaceae and its subfamilies, we performed a fossil-dated analysis of 46 angiosperms, including 38 monocots and 29 grasses. We used age constraints from 11 unambiguous angiosperm fossils (Fig. S3.1) and a secondary calibration on the angiosperm crown node (Silvestro et al., 2015). Including the two old grass fossils and our age estimate for the *Brachypodium* stem node as secondary calibration, versus excluding them, nearly doubled the age for Poaceae (crown node 101–103 Ma vs. 56–71 Ma) and considerably increased the ages for other internal Poaceae lineages, including the PACMAD clade (36–60 vs. 28–41 Ma) and the BOP clade (68–78 vs. 41–50 Ma) (Fig. S3.1, Table 2). The analysis inferred high rate heterogeneity (for cpDNA, the most complete data partition), notably with high substitution rates from around the origin of monocots to the base of Poaceae, with subsequent rate slowdown towards the tips.

### Diversification analyses

We estimated episodic changes in diversification rates in TreePar evaluating two extreme survival probability models (Table 3): for the no extinction scenario (100% of the lineages survive to the next period), the most supported model exhibited two net diversification rate changes within Pooideae at 2 and 30 Ma, although a model with a single rate shift at 2 Ma also received substantial support (Table 3). Under the most supported model, diversification rates are estimated very low (near 0) and turnover high (near 1) during the first time interval. After the first rate-shift at 30 Ma, extinction decreases relative to speciation and diversification increases. After 2 Ma, extinction decreases dramatically and diversification increases again. For the high extinction model (10% survival), the one rate-shift model was selected, with a shift-time at 3 Ma, although a model with a second shift at 37 Ma also received some support (Table 3).

Among the continuous models in RPANDA, a model with diversification varying as a function of the temperature fitted the data better than other constant and time-variable models (Table 4). In the best paleotemperature model speciation increased linearly with the decreasing global temperatures but extinction remained constant (LH= –1313.01, Fig. 1b).

### Phylogenetic signal and ASR of climate niche

Since the diversification analyses revealed an association between changes in temperature and diversification rates, we investigated the ancestral climate conditions of the Pooideae ancestors and their putative exposure to frost and prolonged cold. We used grand total Bioclim means for each species to estimate phylogenetic signal and to reconstruct ancestral states. All estimated Bioclim variables expressed a statistically significant phylogenetic signal, i.e. Pagel’s λ significantly different from zero (*p* <0.001, Table S1.6). Bioclim variables for isothermality (BIO3), temperature seasonality (BIO4) and variables linked to winter season temperatures (BIO6 and BIO11) contained the strongest phylogenetic signals (Pagel’s λ >0.85). For the remaining Bioclim variables Pagel’s λ ranged between 0.30 and 0.85, with variables linked to precipitation containing the weakest phylogenetic signal.

Isothermality (BIO3) was the Bioclim variable with the strongest and most significant phylogenetic signal (Table S1.6). The reconstruction of isothermality as continuous trait (Fig. 2, 3a–d) produced ancestral estimates for the Pooideae ancestor and the Pooideae backbone nodes that were comparable with values of present warm temperate and subtropical climates (Fig. 3a). Isothermality estimates associated with cold temperate and frigid climates were found to have evolved late in the phylogeny and only in a few lineages (Fig. 3b–d).

**Figure 2.**
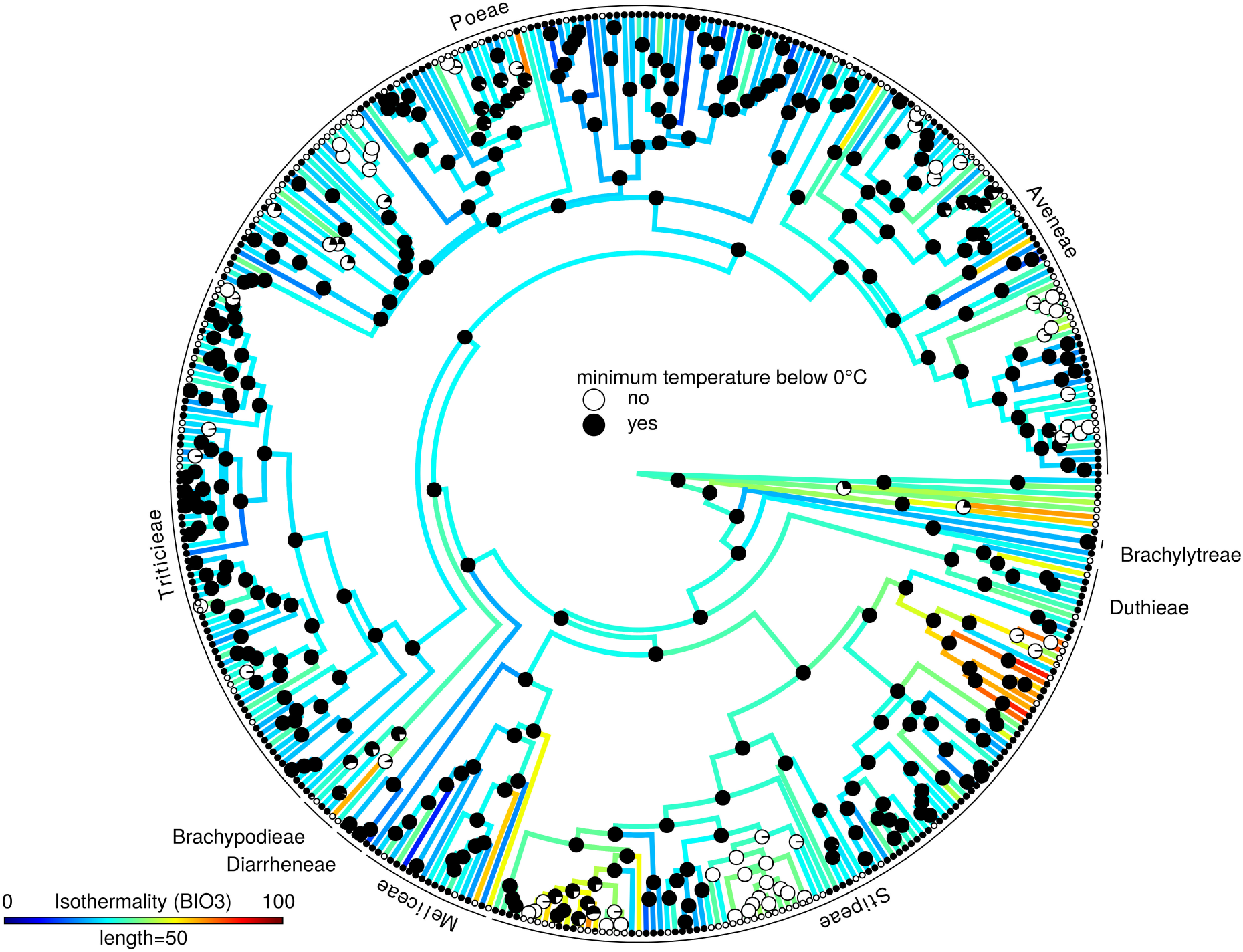
Experience of frost in ancestral Pooideae lineages. The ancestral reconstruction of the minimum temperature of the coldest month (BIO6) as a discrete, binary character (below 0°C yes/no) reveals a high probability of frost-experiencing ancestral Pooideae lineages. The posterior probabilities of the ancestral states are plotted as pie chart diagrams onto the dated phylogeny. Colors on the branches display the reconstructed ancestral values for isothermality (BIO11) shown on the scale bar.

**Figure 3.**
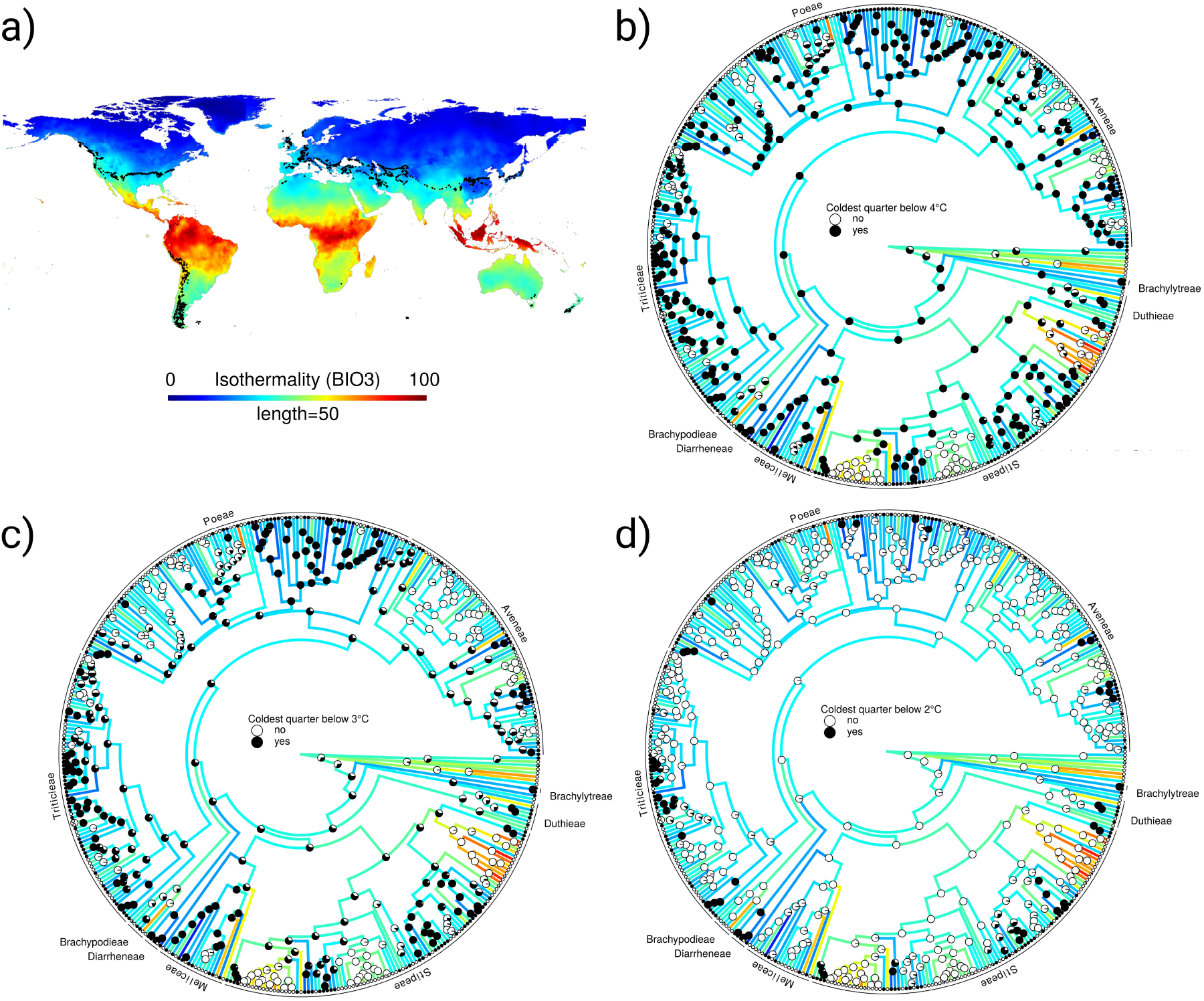
Temperature seasonality and winter severity in ancestral Pooideae lineages. For comparability, global distribution of isothermality values are plotted as color gradient on a contemporary world map a). The scale bar indicates the respective links between color and isothermality value. Black dots in the map indicate coordinates where the mean temperature of the coldest quarter is exactly 2°C. The ancestral reconstructions of the mean temperature of the coldest quarter (BIO11) as three discrete, binary characters (below or not below 4°C, 3°C and 2°C shown in b), c) and d) respectively) reveals that Pooideae ancestors likely experienced mild winters with grand mean temperatures below 4°C, but not below 2°C. The posterior probabilities of the discrete, ancestral states are plotted as pie chart diagrams onto nodes of the dated phylogeny. Branches are colored according to ancestral values for isothermality (BIO3). Colors correspond to scale bar in a).

To elucidate when the Pooideae ancestor adapted to frost and cold winters, we conducted ASR analysis for minimum temperatures of the coldest month (BIO6, Fig. 2), binarized to below vs. above 0°C. This analysis revealed that the ancestor of the Pooideae, as well as the ancestor of each major sublineage, did experience frost (Fig. 2). We scored winter severity as three binary characters derived from the mean temperature of the coldest quarter (BIO11, Fig. 3b–d). Our analyses indicate that ancestors of all major lineages experienced winters with mean temperatures below 4°C (Fig. 3b) and possibly below 3°C (Fig. 3c). However, a mean winter temperature of 2°C is the lower limit for most extant and ancestral lineages (Fig. 3d). Tolerance to such cold winters evolved late in the Pooideae phylogeny in independent lineages.

### Reconstruction of historical biogeography

We reconstructed the historical biogeography of Pooideae performing discrete and continuous ASR analyses. Discrete analyses reconstructed Eurasia as ancestral area of the Pooideae (*pp* = 1), with numerous dispersals into different continents occurring after the divergence of all tribes (Fig. S3.2). Most notably are the independent dispersals of Stipeae lineages to Africa and the Americas. Within Eurasia, the continuous biogeographic analyses centered the origin of the Pooideae MRCA on southwestern Eurasia, at today’s southern part of the Caspian sea (Fig. S3.3). However this estimate is associated with high uncertainty.

## Discussion

### The age of Pooideae

Our age estimates (Fig. 1a) indicate a relatively older age of the Pooideae crown, 66.0 (59.0– 72.6) Ma, than many earlier estimates (Gaut, 2002; GPWG, 2001; Hertweck et al., 2015; Strömberg, 2005) but are in line with recent analyses that have taken into account updated information from the fossil record (Burke et al., 2016; Vanneste et al., 2014; Wang et al., 2015).

Three recently published grass fossils with reliable synapomorphies (Poinar Jr. et al., 2015; Prasad et al., 2005; Wu et al., 2017) strongly contradict previous estimates for young origins of both Poaceae and the internal BOP clade, indicating instead crown ages for Poaceae and subfamily Oryzoideae of at least 101 Ma and 66 Ma, respectively. Including versus excluding the three old calibrations in an angiosperm-wide, four-locus and 46-taxon dataset nearly doubled the age for Poaceae (crown node 101–103 Ma vs. 56–71 Ma) and considerably increased the ages for other internal Poaceae lineages (Fig. S3.1, Tables 2, S1.6).

### Falling temperature throughout the Cenozoic drove diversification in Pooideae

Our analyses suggest cooling temperature as a driver of diversification in Pooideae. Specifically, our paleotemperature-dependent analysis revealed that diversification rates increased as a function of the temperature decrease during the Cenozoic (Fig. 1b). Hence, inferred speciation rates were highest during the coolest intervals of the recent past, notably following the E-O transition, and have generally been increasing since the origin of the Pooideae. The peak in speciation rates toward the present is concurrent with the intensified global cooling trend that culminated with the Pleistocene glaciations. Pooideae might not be unique in this regard. Several radiations following the appearance of temperate biomes have been identified in other plant groups (Favre et al., 2016; Meseguer et al., 2018; Spriggs, Christin, & Edwards, 2014).

A likely driver of the association between cold and diversification is the availability of new niches as the temperate climates greatly expanded across the Holarctic during the mid-late Cenozoic (Eldrett et al., 2009; Liu et al., 2009; Potts & Behrensmeyer, 1992; Pound & Salzmann, 2017; J. Zachos et al., 2001). Preadaptations to endure cold may explain the expansion of Pooideae into temperate climates and increased speciation rates as this lineage successively exploited cooler niches associated with the overall colder climates. Furthermore, higher diversification rates have been found across Poales lineages inhabiting “open” and “dry” habitats compared to lineages inhabiting “shade” and “wet” habitats (Bouchenak-Khelladi, Muasya, & Linder, 2014). For Pooideae, ancestral habitat reconstruction indicates that transitions from “closed” to “open” habitats occurred after the major tribes had diverged (Bouchenak-Khelladi et al., 2010). Thus, increased diversification rates (Fig 1b) not only coincide with climate cooling, but also with transitions to more open habitats. Although our sampling of taxa is relatively low, we would like to stress that it was designed to be exhaustive at the tribe and genus level, which implies that all basal nodes and lineages in the tree have been sampled while the tips are under-sampled. In any case, our RPANDA results are also congruent with the significant increase in diversification rates detected by TreePar in the recent past (Table 3).

### Did the Pooideae ancestor live in cold microhabitats?

Despite the relatively warm global temperatures and the abundance of boreotropical forests at the time of Pooideae origin (Greenwood, Basinger, & Smith, 2010; Pross et al., 2012; Tiffney, 1985; Wolfe, 1975), our reconstruction of the ancestral habitat indicates that ancestors of all major Pooideae lineages experienced and could withstand light frosts and mild winters in a seasonal climate (Fig 2, Figs. 3b–d). Thus, the evidence points to the Pooideae subfamily being a temperate lineage long before the expansion of temperate biomes, contrary to many other temperate plant lineages (Favre et al., 2016; Kerkhoff et al., 2014; Meseguer et al., 2018, 2015). This hypothesis is supported by two recent studies. In a study of three Pooideae species, Zhong, Robbett, Poire, & Preston (2017) identified several gene clusters exhibiting conserved cold response, many of which had previously been characterized as ancient stress response genes. Another study found sixteen cold responsive genes that exhibited conserved expression in five distantly related Pooideae species (Schubert *et al.*, unpublished). Interestingly, most of these genes were induced in response to short-term cold, and are known to be stress-responsive in other angiosperms. Taken together, these results point to a common response to short-term cold stress, but no conserved adaptation to prolonged periods of cold.

The discrete analysis of biogeography identified a Eurasian origin for Pooideae (Fig. S3.2), with numerous dispersals into other continents. This is in line with previous biogeographic analyses of Pooideae (Bouchenak-Khelladi et al., 2010), but with more complete sampling of Pooideae lineages. A continuous biogeographical analysis of the Eurasian taxa centers the origin of Pooideae on southwestern Eurasia (Fig. S3.3). The uncertainty is, however, high and these results must be interpreted cautiously. However, this scenario provides an explanation for the paradoxical early evolution of cold adaptations in a globally warm climate (Fig 2, Fig. 3b–d). We hypothesise that early Pooideae originated in high elevation habitats in the mountains of the nascent Alpine orogeny in southwestern Eurasia. This mountain chain resulted from the collision of the African and Arabian plates with the European (Eurasian) plate from Late Cretaceous onwards, with major phases of mountain building from the Paleocene (Gee & Stephenson, 2006; Moores & Fairbridge, 1997; Sharkov et al., 2015). In cold microhabitats of the nascent Eurasian mountains, the early Pooideae may have evolved some simple stress responses to cold that may have given them sufficient fitness advantage to enable diversification into the temperate niche as temperate climates expanded throughout the Oligocene. The striking delay of some 35 Ma from the origin of Pooideae to the onset of its diversification at the E-O transition (Fig. 1a–b) might reflect long ecological and geographic confinement, e.g. to high-elevation sky islands.

### Lineage-specific adaptations to long winters

Despite the evidence for a temperate ancestral niche (Fig. 2, Figs. 3b–d) our analyses also indicate that tolerance of more extreme temperate conditions, i.e. colder and longer winters, is not shared among the major Pooideae lineages. Coinciding with the intensification of the global cooling trend and the increased seasonality, particularly during and after the E-O transition (Eldrett et al., 2009), we observe emergence of niches with stronger seasonality (low isothermality) and more severe winters (mean temperature of the coldest quarter below 2°C, Fig. 3d) in separate lineages. A similar evolutionary history has been reconstructed for Danthonioideae, where the coldest habitats are occupied by distantly related clades (Humphreys & Linder, 2013). We suggest that complex adaptive pathways for tolerating long, severe winters, (e.g. cold acclimation and adaptations to short growing seasons) evolved independently in Pooideae lineages. Our findings corroborate recent studies of molecular evolution of cold adaptation. Although all species from distantly-related Pooideae lineages are able to cold acclimate, most of the cold responsive genes identified by Schubert *et al.*, (unpublished) were differentially expressed in only one of five investigated species representing different tribes. Furthermore, cold acclimation gene families identified by that study. were mostly exclusive to core Pooideae. Finally, flowering in response to vernalization is widespread in the Pooideae (McKeown, Schubert, Marcussen, Fjellheim, & Preston, 2016). In the core Pooideae, vernalization is highly regulated by the *VRN1* and *VRN2* regulon (Fjellheim et al., 2014). Although cold induction of *VRN1* was found to be an ancestral trait in the Pooideae lineage (McKeown et al., 2016), the regulatory role of *VRN2* is not conserved in the subfamily, but has been co-opted into the vernalization pathway in the core Pooideae (Woods, McKeown, Dong, Preston, & Amasino, 2016).

## Conclusions

Despite an old age of 66.0 (59.0–72.6) Ma, we inferred an ancestrally temperate niche with seasonally occurring frosts for the grass subfamily Pooideae. Although the ultimate sieve for persisting in large parts of the temperate regions is the ability to survive winter, other characters, such as timing of flowering, growth and life history strategies would also have played central roles. Our dated phylogeny forms a rigorous framework for future testing of hypotheses regarding evolution of adaptations to temperate climate from tropical ancestors in light of climate and diversification history of Pooideae. Our study is also the first on grasses to demonstrate the usefulness of employing speciation times, estimated from the entire fossil record of the clade using PyRate, as calibration prior instead of oldest fossil. This method increases objectivity and accuracy in molecular dating, especially for lineages with a sparse fossil record such as grasses, and should be put to use also for other grass lineages where ages are still controversial.

## Data accessibility

All data used in this manuscript are presented in the manuscript and its supplementary material or have been previously published or archived elsewhere.

